# Dropout-based feature selection for scRNASeq

**DOI:** 10.1101/065094

**Authors:** Tallulah S. Andrews, Martin Hemberg

## Abstract

Features selection is a key step in many single-cell RNASeq (scRNASeq) analyses. Feature selection is intended to preserve biologically relevant information while removing genes only subject to technical noise. As it is frequently performed prior to dimensionality reduction, clustering and pseudotime analyses, feature selection can have a major impact on the results. Several different approaches have been proposed for unsupervised feature selection from unprocessed single-cell expression matrices, most based upon identifying highly variable genes in the dataset. We present two methods which take advantage of the prevalence of zeros (dropouts) in scRNASeq data to identify features. We show that dropout-based feature selection outperforms variance-based feature selection for multiple applications of single-cell RNASeq.

## Introduction

Single-cell RNASeq (scRNASeq) has made it possible to analyze the transcriptome from individual cells. In a typical scRNASeq experiment for human or mouse, ~10,000 genes will be detected. Most genes, however, are not relevant for understanding the underlying biological processes, and an important computational challenge is to identify the most relevant features. For some well-studied systems one can find the most important genes by searching the literature, but in most situations an unsupervised approach would be more desirable. However, unsupervised feature selection remains difficult due to the high technical variability and the low detection rates of scRNASeq experiments.

A widely used approach for feature selection is based on the concept of differentially expressed genes, i.e. genes whose level differs between two sets of cells. Differential expression can be considered a supervised method for feature selection, and it is commonly used in bulk RNASeq experiments. However, the conceptual framework is more difficult to apply to scRNASeq since differential expression methods, such as SCDE (Kharchenko et al., 2014), require predetermined, homogeneous subpopulations, which are often unavailable.

Unsupervised methods for identifying relevant features specifically for scRNASeq data have mainly focused on the identification of highly variable genes (Brennecke et al., 2013; Kolodziejczyk et al., 2015; Satija et al., 2015). These methods differ mainly in their approach to adjusting for the relationship between mean and variance inherent to count-data, such as fitting a polynomial regression (Brennecke et al., 2013), binning genes by expression level (Satija et al., 2015), or comparing to a moving median (Kolodziejczyk et al., 2015). Alternatively, specific genes may be identified from their weights inferred during dimensionality reduction (Björklund et al., 2016; Klein et al., 2015; Macosko et al., 2015; Pollen et al., 2014; Usoskin et al., 2015; Wilson et al., 2015). Another approach is to decompose an expression matrix into a small set of meta-features using dimensionality reduction methods, such as principal component analysis (PCA) or t-Distributed Stochastic Neighbor Embedding (t-SNE) (Maaten, Laurens van der and Hinton, 2008). However, these methods are often sensitive to systematic noise, e.g. batch effects, due to the large number of genes subject to technical noise relative to the number of genes influenced by biological effects (Hicks et al., 2015; Tung et al., 2017). In addition, the biological interpretation of extracted meta-features is often challenging.

A salient aspect of scRNASeq data is the presence of “dropouts”, i.e. genes that are not detected in some cells but highly expressed in others. We introduce two novel feature selection methods based on the relationship between dropout-rate and mean expression across genes. The methods are closely related with one specifically adapted for full-length transcripts and the other for UMI-based protocols. Comparing to currently available methods, we find that dropout-rate based methods outperform variance-based methods on both simulated and real datasets.

## Results

### Dropout-based Feature Selection

Dropout-based feature selection is conceptually similar to highly-variable gene-based feature selection. In both cases it is assumed that genes expressed at a constant level will follow some distribution due to technical noise, and that genes responding to a biological perturbation will follow a different distribution (**Figure 1 A,B**). Highly variable gene detection characterizes these distributions using the relationship between the mean and the variance, whereas dropout-based feature selection uses the relationship between the mean and the number of zeros. Since single-cell RNASeq data contains a large number of zeros, with dropout rates often spanning the full range from 0 to 1, this is effective in characterizing the expression distributions (**Table S1**). An important property of the mean-dropouts relation is that differential expression across distinct populations of cells or through pseudotime increases the observed dropout-rate due to the nonlinearity of the relationship (**Figure 1 C**). The advantage of using the dropout-rate over variance is that the former can be estimated more accurately due to much lower sampling noise (**Figure S1**).

**Figure 1:**
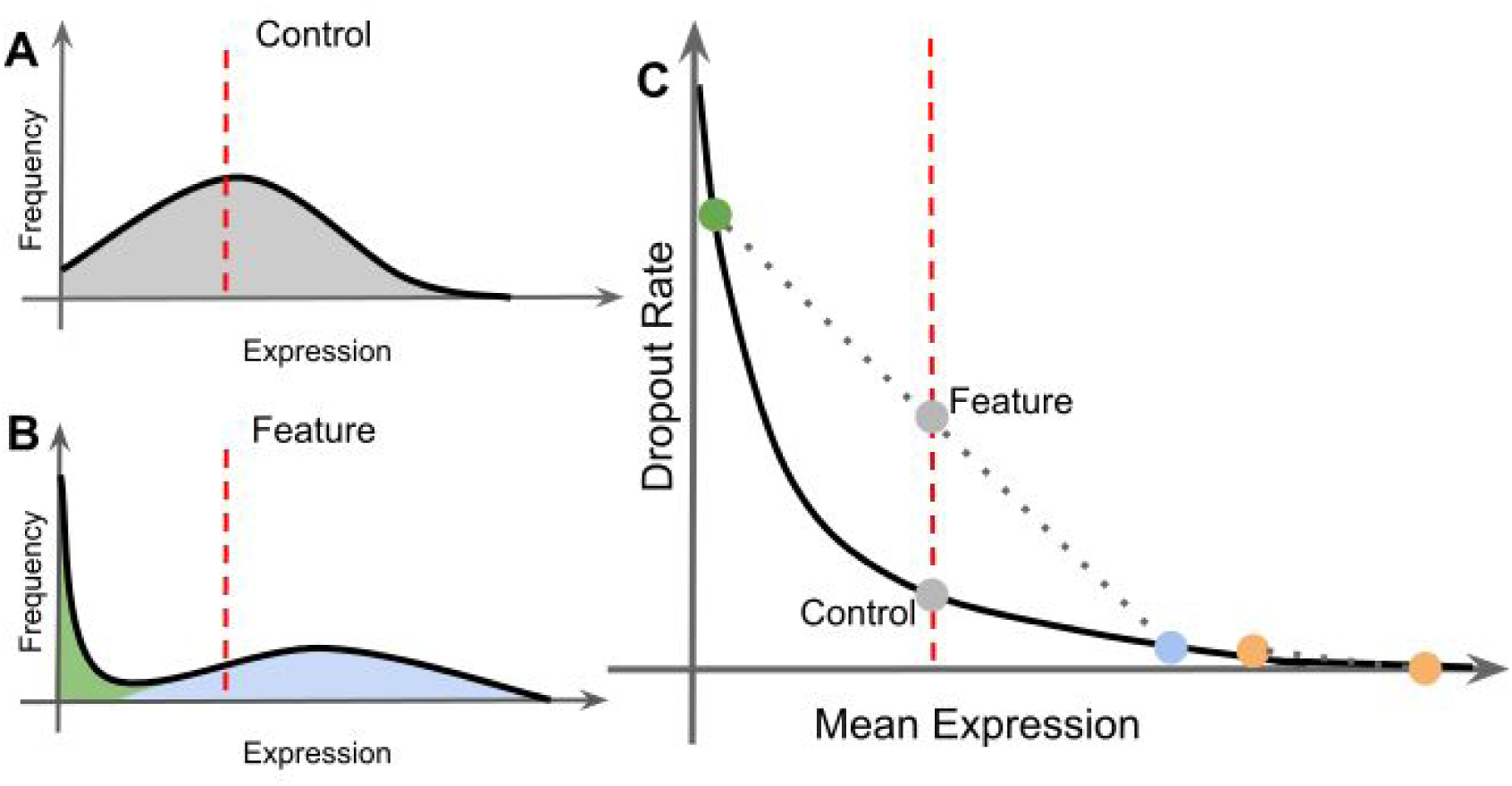
Differentially expressed genes exhibit bimodal expression which increases the dropout rate relative to the mean expression. (**A & B**) Genes with the same mean expression (dashed red line), but (A) is expressed evenly across cells, whereas (B) is highly expressed in some cells (blue) and lowly expressed in others (green). (**C**)This leads to a surplus of dropouts since mean and dropout rate average linearly (dotted line) whereas the expectation (black line) is non-linear. Orange points indicate a gene with very high expression where differential expression leads to only a small increase in dropout-rate.

The first novel method for identifying high-dropout genes fits a Michaelis-Menten function to the relationship between mean expression (*S*) and dropout-rate (**M3Drop**).

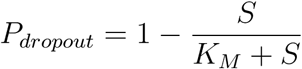

Since the Michaelis-Menten function has a single parameter (K_M_), we can test the hypothesis that the gene-specific K_i_ is equal to the K_M_ that was fit for the whole transcriptome. This can be done by propagating errors on both observed dropout rate and observed mean expression to estimate the error of each K_i_. The significance can then be evaluated using a t-test (see: Methods). We confirmed that the M3Drop model fits three diverse Smartseq/2 scRNASeq datasets (**Figure S3d-f**).

The second method for identifying high dropout genes (**NBDrop**) was tailored for low-depth high-throughput data, which generally employs unique molecular identifiers (UMIs). NBDrop fits a library-size adjusted negative binomial model to account for the strong dependency between library-size and number of detected genes for high-throughput protocols (**Figure S2**)(Grün et al., 2014). We use the mean (μ_ij_) and dispersion (r_j_) parameterization of the negative binomial distribution for the number of transcripts *Y*:

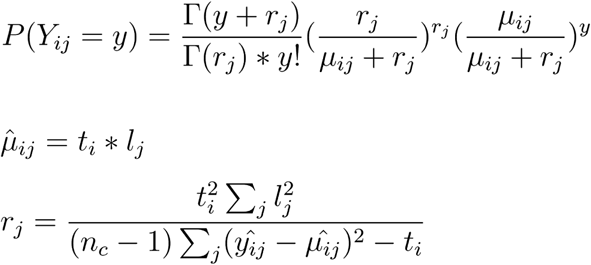

where t_i_ is the total counts for gene i, l_j_ is the relative library size of cell j, n_c_ is the total number of cells, and 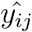 are the observed counts of gene i in cell j.

To calculate the expected dropout-rate, we first fit a log-linear relationship between the average expression 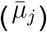 and dispersion (r_j_) across all genes. This is used to estimate the gene-specific dispersions under the null hypothesis (r_0j_), which is then used to calculate expected dropout rate (p_j_):

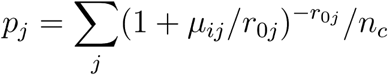

Features with observed dropout-rate, *d_j_*, that is significantly higher than expected by chance were identified using a binomial test. As a null hypothesis, it is assumed that the dropout-rate follows a binomial distribution, d_j_ ~ Binom(n_c_, p_j_), and we use this to identify genes where d_j_ is significantly higher than expected by chance.

Unlike M3Drop, NBDrop does not account for errors in the estimated mean expression levels. Thus, it is not as well suited for data with small-sample sizes and/or high amplification noise, as is typical of full-transcript, plate-based protocols (Islam et al., 2014). We confirmed that the NBDrop model fits three diverse tag-based scRNASeq datasets (**Figure S3a-c**).

### Comparison to Other Feature Selection Methods

To demonstrate the advantage of using dropouts for feature selection, we compared both M3Drop and NBDrop to their variance-based counterparts. For comparison with NBDrop, we used the same library-size adjusted negative binomial model but ranked genes by the distance between their gene-specific dispersion and the expected dispersion calculated from a linear relationship fit to the log-transformed means and dispersions of across all genes (Methods).

For comparison with M3Drop we considered the popular highly variable gene (**HVG**) method proposed by Brennecke et al. (2013) (Brennecke et al., 2013). HVG fits a quadratic model to the relationship between mean expression and the coefficient of variation squared (CV^2^) and outliers above the fitted curve are selected as features. Originally, the model was fit using ERCC spike-ins, but this can be problematic since many datasets do not contain spike-ins or may have technical issues which affect the consistency of spike-ins added to all experimental batches (Svensson et al., 2017). Thus for consistency we fit the model using all genes, based on the assumption that most genes exhibit only technical noise.

In addition, we considered feature selection based on the Gini Index (Gini, 1912) a common measure of skewness. We use the method proposed in GiniClust (Jiang et al., 2016) to account for the relationship between the Gini Index and the maximum observed expression and estimate a p-value for each gene (**Gini**).

As an alternative to the above methods which all rely on identifying outlier genes based summary statistics, we considered gene-gene correlations (**Cor**), which ranked genes based on the magnitude of their strongest correlation, and principal component analysis (**PCA**), which ranked genes by their loadings for the top PCs calculate for the whole dataset. Finally we considered a consensus approach (**Cons**) where genes were ranked by each method independently then the average rank across all methods was used to score each feature.

### Evaluating Feature Selection Methods Using Synthetic Data

In general, the main goal of unsupervised feature selection in scRNASeq analyses is to identify genes that are differentially expressed according to some unknown underlying structure in the data. Thus, we evaluate the ability of feature selection methods to identify genes that are differentially expressed between two populations of cells from scRNASeq data when the labels for the two groups are not provided.

We simulated data using both a zero-inflated negative binomial model (ZINB) and a library-size adjusted negative binomial model (LS-NB). The parameters of the generative models were set by fitting to three different full-transcript and three different tag-based scRNASeq datasets (Methods). Data were simulated for two cell populations where the minor population constituted between 1–50% of the total number of cells, generating a total of 108 simulated datasets. Log-fold changes in mean expression for each gene was drawn from a normal distribution, and genes with a log-fold change of >5 between the cell populations were considered true DE genes and genes with log-fold change of < 1 were considered as not DE. For a fair comparison of methods we rank all genes by significance or effect-size for each method and calculate the area under the ROC curve (AUC) based on these rankings.

Both dropout based feature selection methods, NBDrop and M3Drop, performed significantly better than variance-based feature selection on the ZINB simulations (**Figure 2**). Furthermore, NBDrop significantly outperformed all other outlier-detection based methods on LS-NB simulations. Notably, the popular HVG method was only marginally better than random chance (AUC < 0.6) when we considered only the top 2000 ranked features for both sets of simulations. The best method was gene-gene correlations, which slightly outperformed the dropout-based methods. In addition to considering the ranking of the genes, we also considered the number of genes reported at a 5% FDR. Here we found big differences between the methods since only NBDrop, M3Drop and HVG have reasonable estimates of feature significance (**Figure 2 E**). The Gini index detects almost no significant features and gene-gene correlations detects more than 80% of genes as significant features, while PCA and NBDisp do not have significance tests associated with them. In addition, gene-gene correlations took hours to compute vs seconds for the outlier-based methods. Greater inequality in relative size of the two populations of cells decreased the performance of all feature selection methods except PCA (**Figure S4**).

**Figure 2:**
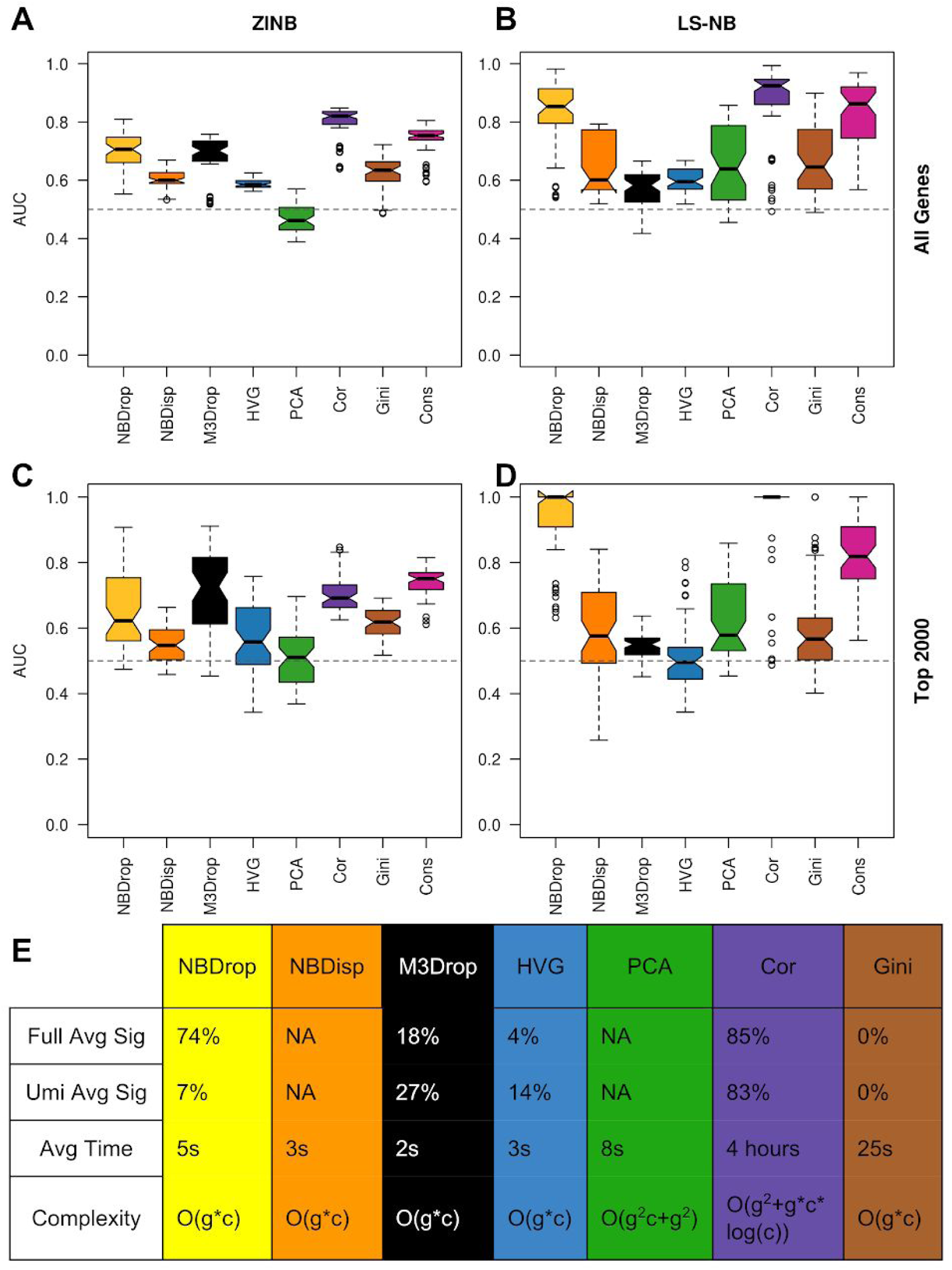
Feature selection performance on simulated data. Each feature selection method was applied to 54 ZINB (**A, C**) and 54 LS-NB (**B, D**) simulated datasets. Boxplots indicate the area under the receiver operator curve (AUC) for identifying true DE genes when all genes (**A, B**) or only the top 2000 features (**C, D**) were considered. Notches indicate 95% confidence intervals of the median. Dashed line indicates an AUC of 0.5 which is expected by chance. (**E**)

Proportion of genes identified as significant features after applying a 5% FDR correction to the p-values reported by each feature selection method when applied to the simulated datasets. Theoretical computational complexity with respect to number of genes (g) and number of cells (c), and observed runtimes (CPU time).

One potential disadvantage of dropout-based feature selection is that they may be unable to detect highly expressed genes since these may have no dropouts, even when they are differentially expressed across cell populations. However, when we binned data by expression level we found that dropout-based feature selection performance only dropped below variance-based feature selection for the top 5% most highly expressed genes in our simulations. This corresponds to a mean expression level of >1,000 reads/cell or >64 umis/cell (**Figure S5**), levels that are found only rarely in most datasets. For the other 95% of genes, dropout-based feature selection out-performed variance-based feature selection. Dropout methods perform better in part because estimates of variance are very sensitive to sampling noise, particularly for highly skewed distributions (Rose and Smith, 2002), including those that are frequently observed for single-cell RNASeq datasets. Our simulations show that sampling noise can produce substantial errors even in datasets of 1,000 cells (**Figure S1**). In addition, highly variable gene and high dropout feature selection scale more favorably with the number of cells than gene-gene correlations (**Table S2**).

### Feature Selection for Real Datasets

To account for other sources of technical noise that were not accounted for in our simulations, we tested the feature selection methods on two real scRNASeq datasets. Again, we considered a scenario where the two groups of cells are assumed to be unknown. We selected the Tung et al., (2017) human iPSCs as an example of tag-based data and the Kolodziejczyk et al., (2015) mouse ESCs as an example of full-transcript data. In both studies, the authors examined homogeneous cultures and they performed multiple replicates of bulk RNASeq on at least two of the same conditions at were used for scRNASeq. We use the differentially expressed genes identified from the bulk RNASeq data as ground truth to test the performance of feature selection on the single-cell data.

Again, both dropout-based feature selection methods significantly outperformed variance-based methods on full-transcript datasets recapitulating our results on simulated data (**Figure 3 top**). On the Tung data NBDrop significantly outperformed all other methods except the consensus. Interestingly, on both real datasets the consensus features significantly outperformed all other methods. One reason for this may be that correlation and PCA based feature selection pick up complementary information from the outlier-detection based feature selection (**Figure 3 bottom**) and different biases with respect to mean expression level (**Figure S6**)

**Figure 3:**
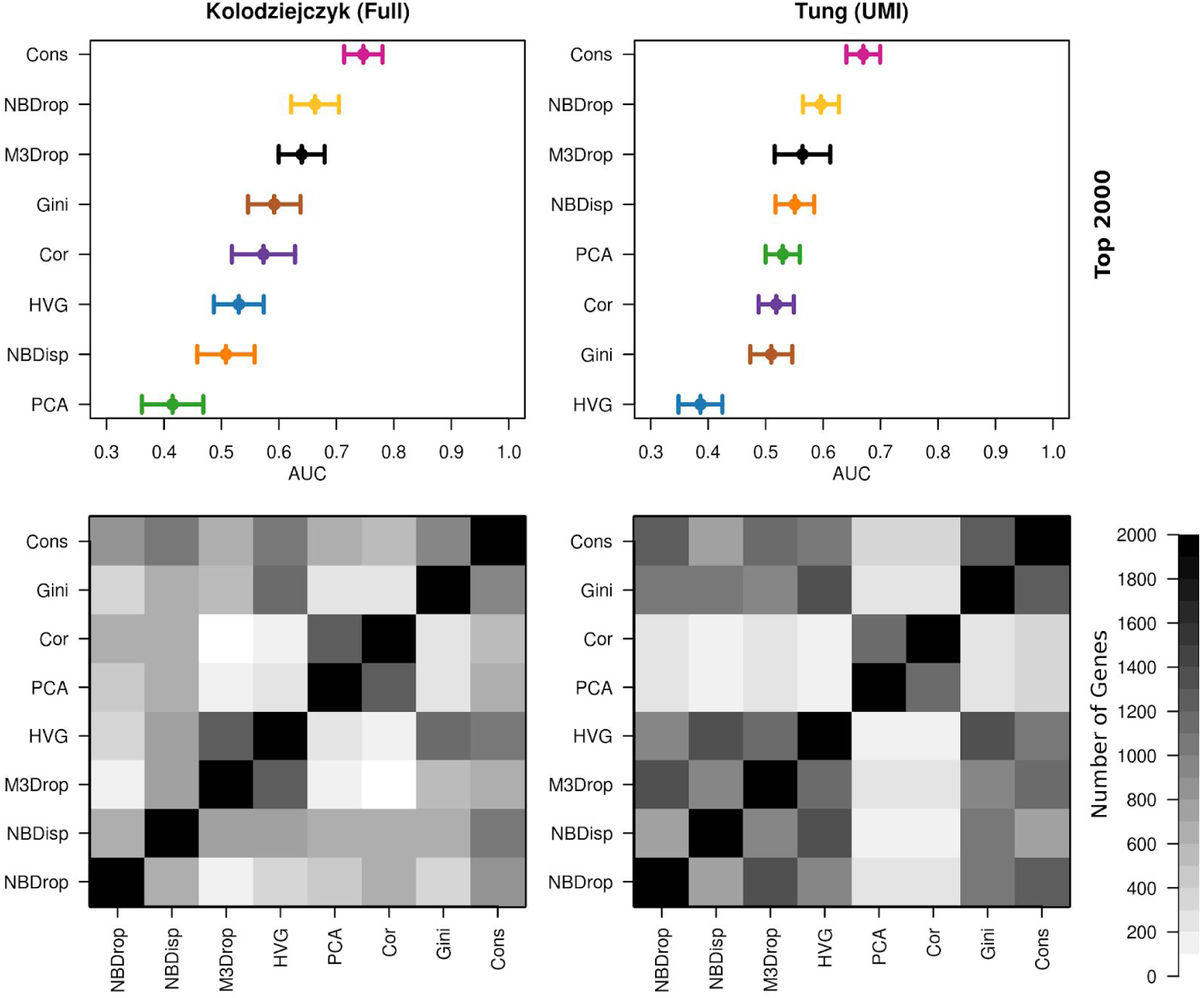
Performance of feature selection on real scRNASeq data. (top row) performance of top 2000 features as ranked by each method on real single-cell RNASeq data. Consusens DE genes from bulk were used as the ground truth. Error bars indicate 95% CIs. We considered both a Smartseq2 dataset (Kolodziejczyk, left column) and a UMI-tagged (Tung, right column) dataset. (bottom row) Overlaps among the top 2000 features identified by each method, Cor and PCA methods are distinct because they are biased towards more highly expressed genes (**Figure S6**).

### Reproducibility of Selected Features

Combining different scRNASeq datasets collected from different labs and using different protocols remains a challenging effort due to the prevalence of batch effects (Haghverdi et al., 2017). To demonstrate how unsupervised feature selection can be used to identify genes that are biologically informative across multiple scRNASeq datasets, we considered five datasets examining early mouse embryo development (**Table S1**). Since early development is a synchronized, robust biological process, we can use the sampling time-point of each cell as ground truth (Kiselev et al., 2017). We evaluated how reproducible features were across datasets, and how well they capture both the distinct stages and developmental trajectory as opposed to the batch-effects between datasets (**Figure 4**). For each feature selection method, we extracted the top 2,000 features from each dataset, and we then asked how well the gene lists agreed. Gene-gene correlations, both dropout-based feature selection methods and the consensus features were most reproducible with ~5% of the top 2000 features detected in all five datasets (**Figure 4A**). To further investigate the biological relevance of the features identified in three or more of the datasets by each method, we used hierarchical clustering to group the data using only the set of features that were reproducible in at least three datasets. (**Figure 4B**). Comparing the clusters with the ground truth developmental stages, we conclude that the dropout-based features produced the most accurate clustering whereas the PCA and gene-gene correlation features were least accurate. In addition, we considered whether the ordering of the stages through development was preserved by estimating pseudotime with DPT (Haghverdi et al., 2016). Again, we found that dropout-based features performed the best, whereas gene-gene correlation and particularly PCA features performed the worst. Thus, the dropout-based feature selection methods were most accurate in identifying biologically relevant genes, consistent with the findings in our related paper on a cross-dataset comparison method (Kiselev and Hemberg, 2017).

**Figure 4:**
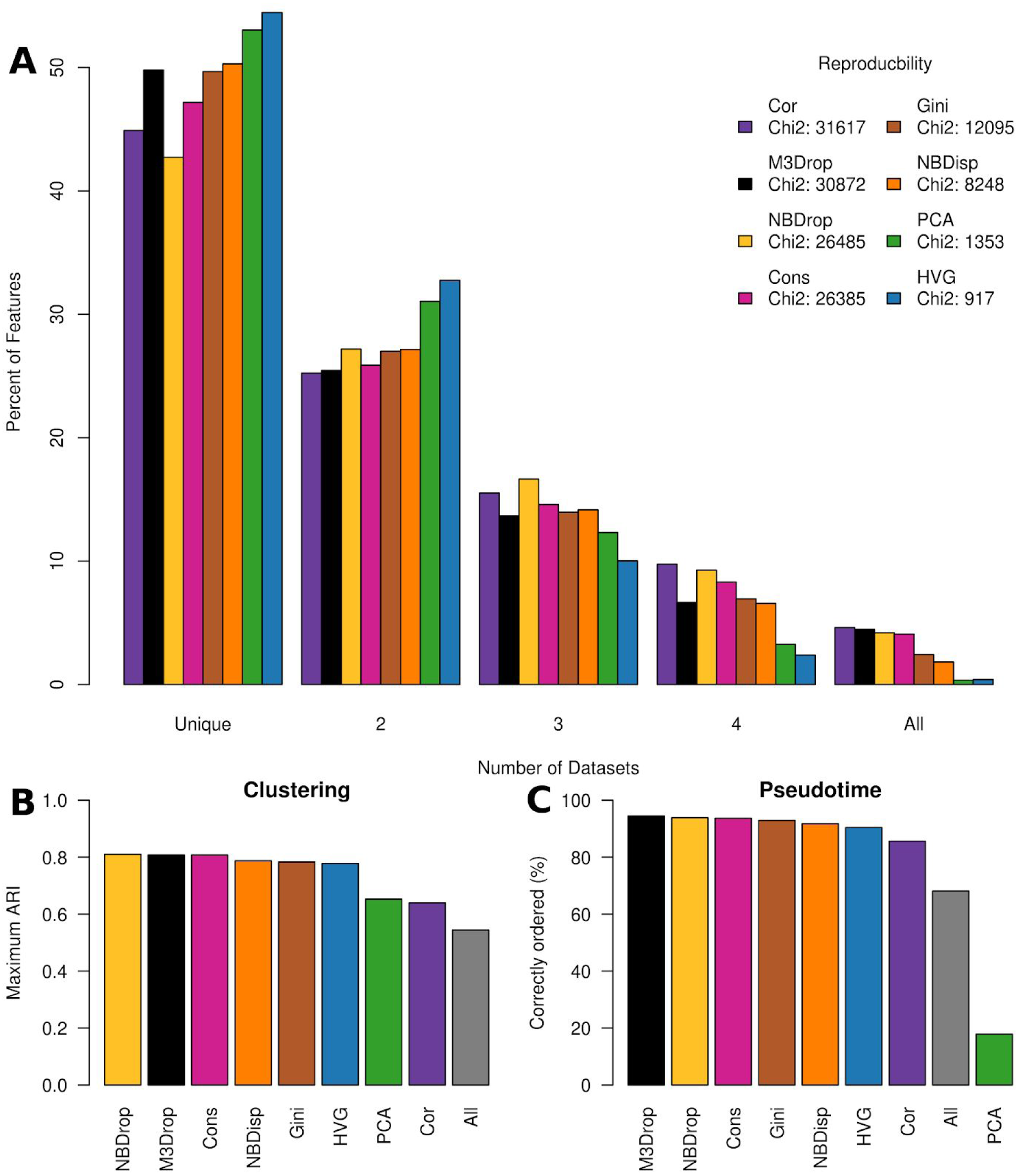
Biological relevance of identified features. (**A**) Reproducibility of features across five early mouse development datasets. The top 2000 features identified by each method in each dataset were intersected with each other. To summarize the extent of overlap we computed the *X*^2^ statistic: Σ(obs-exp)^2^/exp, where the sum is over the number of datasets (see Methods). (**B**) Accuracy of clustering the combined datasets compared to the true developmental stages using the adjusted Rand index (ARI). The value shown is the ARI after Ward hierarchical clustering using just those genes identified in at least three of the five datasets. (**C**) Percentage of cells in the correct order (zygote->blastocyst) after inferring pseudotime using DPT (Haghverdi et al., 2016) on the combined datasets with just those genes identified in at least three of the five datasets.

## Discussion

Due to the high level of noise affecting all genes in scRNASeq experiments, feature selection is an important step in the data analysis. We present two new method for feature selection in single-cell RNASeq data based on identifying high dropout-genes. M3Drop uses a Michaelis-Menten equation to model the zeros in full-transcript scRNASeq. The other model, NBDrop, uses a modified negative binomial model to account for differing sequencing depths or tagging efficiency of UMI-tagged scRNASeq data. Feature selection using these models outperforms existing methods on both real and simulated datasets (**Figure 2, 3**). Furthermore, we show that the dropout-based feature selection can help overcome batch-effects between datasets to reveal underlying biological processes (**Figure 4**).

Here we have considered several different feature selection methods and evaluated them on both simulated and real scRNASeq data. In simulated data, with no technical confounders, gene-gene correlations out-perform all other methods of feature selection (**Figure 2**). However, in real datasets where other confounders such as stress, RNA degradation, or reagent batch can create spurious gene-gene correlations we found that the correlation-based method performed similarly to variance-based feature selection methods and was out-performed by our dropout-based approaches (**Figure 3, 4**). Despite the similar overall performance, correlation and PCA based feature selection identified distinct features from those identified by high-variance or high-dropout based methods (**Figure 2 B**). As a result, taking the consensus of several methods outperformed any individual method.

We have demonstrated that dropout-based unsupervised feature selection can identify biologically meaningful genes. For both M3Drop and NBDrop, the features can be interpreted as differentially expressed genes, but since the methods do not require subpopulations to be defined *a priori*, they are widely applicable. Interestingly, we found differences in performance on different types of data, for instance the popular HVG detection method (Brennecke et al., 2013) has poorer performance on UMI-tagged data than on full-transcript data (**Figure 2,3**) thus highlighting the importance of tailoring models to account for differences between protocols. As a result we develop two methods of dropout-based feature selection, M3Drop tailored to Smartseq/2 platforms and NBDrop tailored to UMI-based platforms. We expect that this effect is not limited to feature selection, but that the choice of protocol will affect many statistical analyses of scRNASeq data. Thus it is important to consider the type of data each method was designed to handle when choosing an analysis pipeline for scRNASeq datasets.

## Methods

### Dropout-based feature selection methods

#### M3Drop

The expression of each gene was averaged across all cells including those with zero reads for a particular gene (*S*). Dropout rate was calculated as the proportion of cells with zero reads for that gene (*P_dropout_*). We fit the Michaelis-Menten equation (Michaelis and Menten, 1913) to the relationship between these two variables

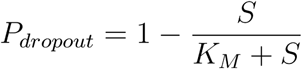

using maximum likelihood estimation as implemented by the mle2 function in the bbmle R package to obtain the global *K_M_* across all genes. This model fits full-transcript data very well (**Figure S2 D-F**). The Michaelis-Menten equation can be rearranged with K on the left hand side. This is useful for estimating a gene-specific *K_j_* as

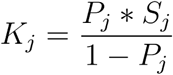

Since *K_j_* is a function of both the dropout and the mean expression, the measurement error for each *K_j_* estimate was calculated using error propagation rules to combine errors on observed S and P:

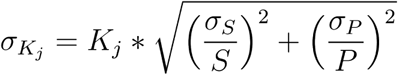

Where *σ_S_* is the sample standard deviation of S and *σ_Ρ_* is the sample standard deviation of P. The K_j_’s were observed to be log-normally distributed around the globally fit K_M_ (**Figure S6**). Thus, we tested each one against the global K_M_ that was fit to the entire dataset using a one-sided Z-test:

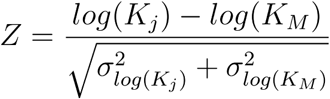

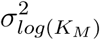 was estimated as the standard error of the residuals and added to 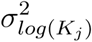

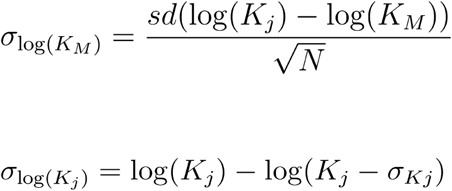

The resulting p-values can be used to ascertain the significance of each feature or to rank genes in order of decreasing significance (increasing p-value). Since all computations are based on gene-level statistics the method scales linearly with the number of cells and number of genes.

#### NBDrop & NBDisp

Negative binomial models have been shown to fit normalized molecule counts from single-cell RNASeq data employing unique molecular identifiers, referred to as UMI-tagged data (Grün et al., 2014; Islam et al., 2014). We modified the single negative binomial distribution to explicitly model the tagging/sequencing efficiency for each cell (*l_j_*) as the relative total molecules observed in cell *j*.

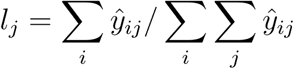

where 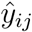 is the number of UMIs observed for gene *i* in cell *j*. Thus, each observation is modelled as a negative binomial model with mean and variance equal to:

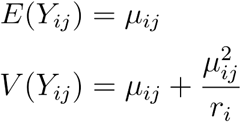

The mean (*μ*_ij_) and the gene-specific dispersion parameter (r_i_) are estimated as:

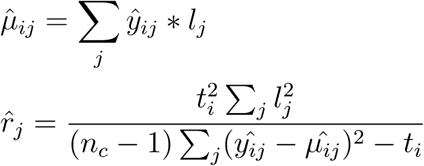

where *t_i_* is the total molecules counts for gene *i*, and *n_c_* is the total number of cells. Genes with Poissonian behavior, which results in negative dispersion, were assigned a maximum dispersion of 10^10^. To identify high dispersion genes using this model (**NBDisp**) we fit a linear regression between the log observed mean expression and the log estimated dispersion parameter across all genes with mean expression > 16 and non-Poissonian dispersions. The residuals from this regression was used to rank genes.

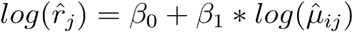

The probability of any given observation being a zero (dropout) is calculated as:

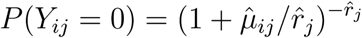

And the expected total dropouts per gene is:

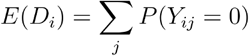

To identify high dropout genes we substituted the expected dispersion calculated from β_0_ and β_1_ in the linear regression equation above for the gene-specific dispersions, and model the number of dropouts per gene as a binomial distribution (**Figure S2 A-C**). The p-value of the observed dropouts can be used to test significance of features or to rank them. This method also scales linearly in number of genes and cells.

### Accuracy using bulk RNASeq as ground truth

To evaluate the accuracy of feature selection methods we use two published datasets for which bulk RNASeq data was available in addition to single-cell data. For both datasets, the cell populations are relatively homogeneous. Tung et al., (2017) considered iPSCs from three different individuals and performed three replicates of UMI-tagged scRNASeq and three replicates of bulk RNASeq for each. Read/UMI counts were obtained from the Gene Expression Omnibus (GSE77288).

For Kolodziejczyk et al., (2015) we considered ESCs grown under two conditions: alternative 2i and serum for which there were three replicates of scRNASeq and two replicates of bulk RNASeq. Full-transcript single-cell RNASeq read counts were obtained from ArrayExpress (E-MTAB-2600). Raw reads for bulk samples were obtained from ArrayExpress and mapped to GRCm38 using STAR (Dobin et al., 2013) and gene level read counts were obtained using featureCounts (Liao et al., 2014).

The ground truth differentially expressed genes were obtained for each dataset using three standard analysis methods: DESeq2 (Love et al., 2014), edgeR (Anders et al., 2013), and limma-voom (Law et al., 2014). Genes identified as differentially expressed using all three methods at 5% FDR were considered ground-truth positives. Genes not identified as DE by any of the methods at 20% FDR were considered ground-truth negatives. This resulted in 1,915 positives, and 8,398 negatives for the iPSCs; and 709 positives and 11,278 negatives for the ESCs (**Figure S7**).

Accuracy of feature selection methods was evaluated using the area under the receiver operator curve (AUC) using the pROC R package, and the number of DE genes among the top 2000 ranked genes.

### Single-cell RNASeq datasets

We considered eleven public scRNASeq datasets (**Table S2**). These were chosen to reflect a range of different dataset sizes, sequencing methods and cell-types. Datasets where the expression matrix consisted of raw read counts (or UMI counts) were normalized using counts per million except for NBDrop and NBDisp. Quality control was performed prior to all analyses as follows. First, all genes annotated as processed pseudogenes in Ensembl (version 80) were removed and cells with fewer than 2000 detected genes were removed. Genes detected in fewer than 4 cells or with average normalized expression < 10^−5^ were excluded from consideration. For the Deng data, only single mouse embryo cells analyzed using the SmartSeq protocol were considered to avoid technical artefacts.

### Simulated datasets

We simulated UMI-tagged data using the depth-adjusted negative binomial model fit to one of the three UMI-tagged datasets, Tung (Tung et al., 2017), Zeisel (Zeisel et al., 2015) and Klein (Klein et al., 2015). Mean gene expression levels were drawn from a log-normal distribution fit to the respective dataset. Cell-specific sequencing efficiency was drawn from a gamma distribution. Finally, gene-specific dispersions were calculated from the mean expression level using the power-law relationship fit to the respective dataset. Each simulated dataset contained 25,000 genes and 500 cells and were consistent with the original data (**Figure S8,S9**).

We simulated full-transcript data using a zero-inflated negative binomial model fit to each of three full-transcript datasets, Pollen (Pollen et al., 2014), Buettner (Buettner et al., 2015), or Kolodziejczyk (Kolodziejczyk et al., 2015). As before, mean gene expression levels were drawn from a log-normal distribution and gene-specific dispersions were calculated from the mean expression level using a power-law relationship. Simulated expression values were inflated with zeros according mean expression using the Michaelis-Menten equation fit to the respective dataset. Since full-transcript data is generally obtained from fewer cells than UMI-tagged data, each simulated full-transcript dataset contained 200 cells and 25,000 genes and were consistent with the original data (**Figure S8,S9**).

Differentially expressed (DE) genes were added by increasing/decreasing the mean expression of each gene in a subset of the cells by a log base-2 fold change drawn from a normal distribution with mean = 0 and sd = 2. Dispersions were adjusted in the differentially expressing cells according to the fitted power-law relationship. We considered subpopulations containing 1%, 10%, 20%, 30%, 40% or 50% of the cells and three replicates for each of the six dataset and each subpopulation size were generated.

Genes with a greater than 5-fold increase or decrease in mean expression were considered ground truth DE genes respectively. Genes with less than absolute 1-fold change in mean or dispersion were considered unchanged.

### Analysis of Developmental Datasets

#### Reproducibility

We considered five full-transcript single-cell RNASeq datasets examining mouse embryonic development from fertilization to blastocyst (**Table S2**). Any genes not present in all five datasets were removed, leaving 11,315 genes in total and the top 2000 ranked genes for each feature selection method were compared across datasets. The expected number of genes identified in precisely *n* datasets was calculated as:

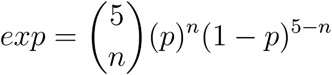

where *p* is the proportion of genes identified by each method i.e. *p* = 2,000/11,285 = 0.18. This was summarized across all *n* using a chi-square statistic:

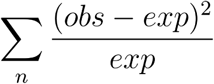

Biological analyses were performed on the subsets of features that were identified in at least three of the five developmental datasets by each method respectively. Expression levels were converted to log2 counts-per-million prior to analysis. Known biological stages were collapsed to “oocyte/zygote”, “2-cell”, “4-cell”, “8-cell”, “16-cell/morula”, and “blastocyst” since there is little evidence to support biologically meaningful differences at different timepoints within these groups.

Clustering was performed using Ward’s hierarchical clustering (Ward, 1963) on Euclidean cell-cell distances. The resulting tree was cut at every possible height and the maximum adjusted-Rand-index (Rand, 1971) between the clusters and the known stage was reported.

Pseudotime was estimated using the DPT/destiny R-package. Cells were placed in order and in reverse order according to the inferred pseudotime, and their respective stage labels were compared to the “ideal” ordering, i.e. all zygotes then all 2-cells etc…, the proportion of matches between the two sets of labels was used as the percent of cells correctly ordered. The ordering, forward or reverse, which gave the highest percent correct was reported.

### Code/Data Availability

Dataset accession codes are listed in **Table 1**.

R packages used: ROCR (version 1.0–7), gplots (version 3.0.0), bbmle (version 1.0.18), reldist (version 1.6–6), irlba (version 2.1.2), monocle (version 2.2.0), destiny (version 1.0.0), DESeq2 (version 1.10.1), edgeR (version 3.12.1), limma (version 3.26.9)

M3Drop and NBDrop & NBDisp are freely available on github (contains code for highly variably gene detection): https://github.com/tallulandrews/M3Drop

## Author Contributions

TA and MH conceived of the project and wrote the manuscript. TA developed the method, produced the code, analyzed and interpreted the data. MH supervised the work.

## Acknowledgements

The authors would like to thank: Vladimir Kiselev, Davis McCarthy, Simon Andrews, and Tomislav Ilicic for their comments and suggestions for improving this manuscript.

## Supplementary Tables

**Table S1:**
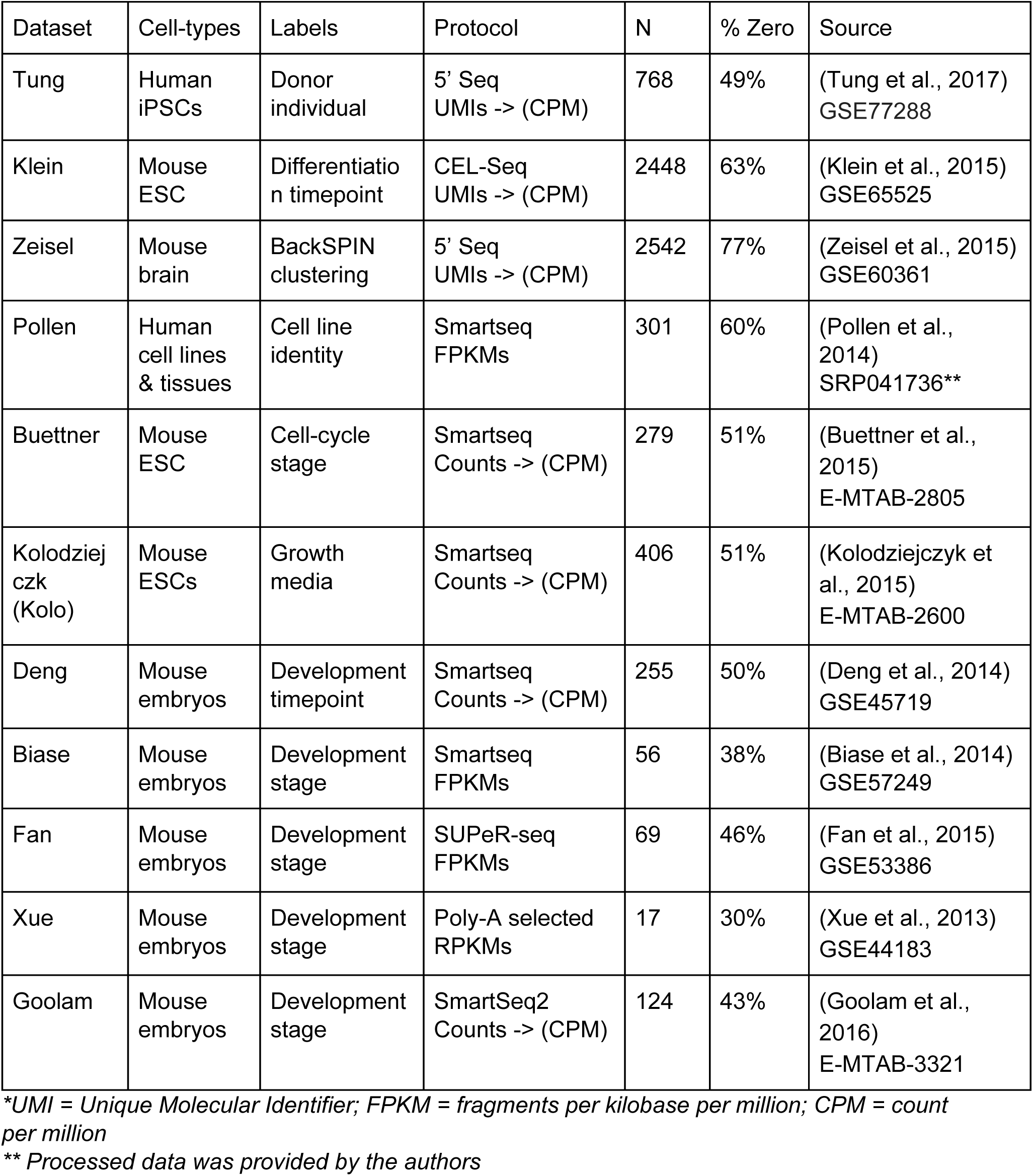
Publicly available single-cell RNASeq datasets, normalization method and proportion of zero valued entries in the filtered expression matrix.

| Dataset | Cell-types | Labels | Protocol | N | % Zero | Source |
| --- | --- | --- | --- | --- | --- | --- |
| Tung | Human iPSCs | Donor individual | 5' Seq UMIs -> (CPM) | 768 | 49% | (Tung et al., 2017) GSE77288 |
| Klein | Mouse ESC | Differentiation timepoint | CEL-Seq UMIs -> (CPM) | 2448 | 63% | (Klein et al., 2015) GSE65525 |
| Zeisel | Mouse brain | BackSPIN clustering | 5' Seq UMIs -> (CPM) | 2542 | 77% | (Zeisel et al., 2015) GSE60361 |
| Pollen | Human cell lines & tissues | Cell line identity | Smartseq FPKMs | 301 | 60% | (Pollen et al., 2014) SRP041736** |
| Buettner | Mouse ESC | Cell-cycle stage | Smartseq Counts -> (CPM) | 279 | 51% | (Buettner et al., 2015) E-MTAB-2805 |
| Kolodziejczk (Kolo) | Mouse ESCs | Growth media | Smartseq Counts -> (CPM) | 406 | 51% | (Kolodziejczyk et al., 2015) E-MTAB-2600 |
| Deng | Mouse embryos | Development timepoint | Smartseq Counts -> (CPM) | 255 | 50% | (Deng et al., 2014) GSE45719 |
| Biase | Mouse embryos | Development stage | Smartseq FPKMs | 56 | 38% | (Biase et al., 2014) GSE57249 |
| Fan | Mouse embryos | Development stage | SUPeR-seq FPKMs | 69 | 46% | (Fan et al., 2015) GSE53386 |
| Xue | Mouse embryos | Development stage | Poly-A selected RPKMs | 17 | 30% | (Xue et al., 2013) GSE44183 |
| Goolam | Mouse embryos | Development stage | SmartSeq2 Counts -> (CPM) | 124 | 43% | (Goolam et al., 2016) E-MTAB-3321 |

\*UMI = Unique Molecular Identifier; FPKM = fragments per kilobase per million; CPM = count per million
\*\* Processed data was provided by the authors

## Supplementary Figures

**Figure S1.**
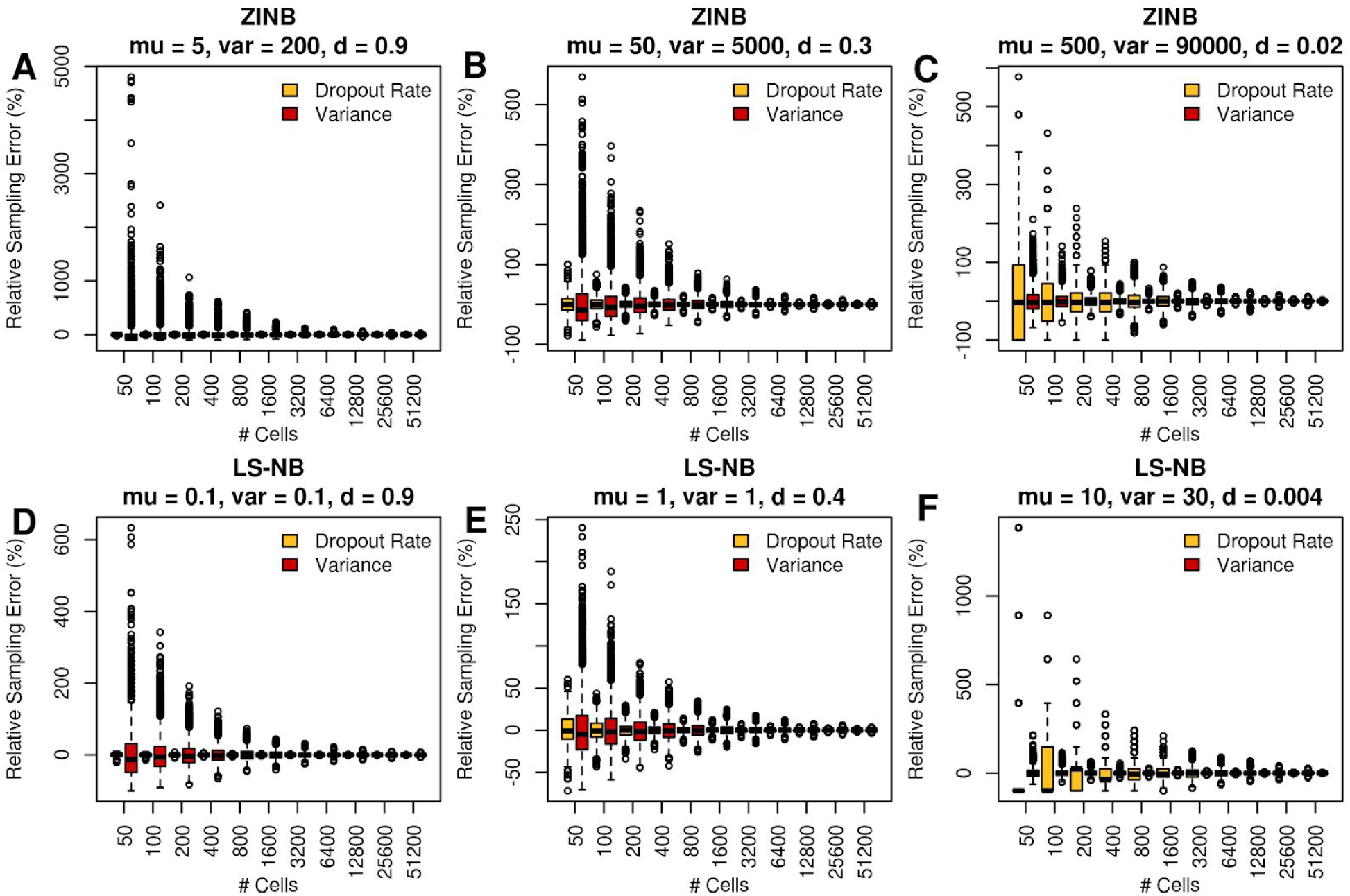
Sampling noise for sample variance and sample dropout rate in a homogeneous population. Simulated expression values for 1,000,000 cells were downsampled without replacement 10,000 times. The percent error of the observed variance/dropout rate of the downsampled values relative to the variance/dropout rate calculated over all 1,000,000 values was calculated. (**A-C**) Simulating from a zero-inflated negative binomial fit to Smartseq2 data. (**D-F**) Simulating from the library-size adjusted negative binomial fit to UMI-tagged data. mu = mean, var = variance, d = dropout-rate for all 1 million simulated cells.

**Figure S2:**
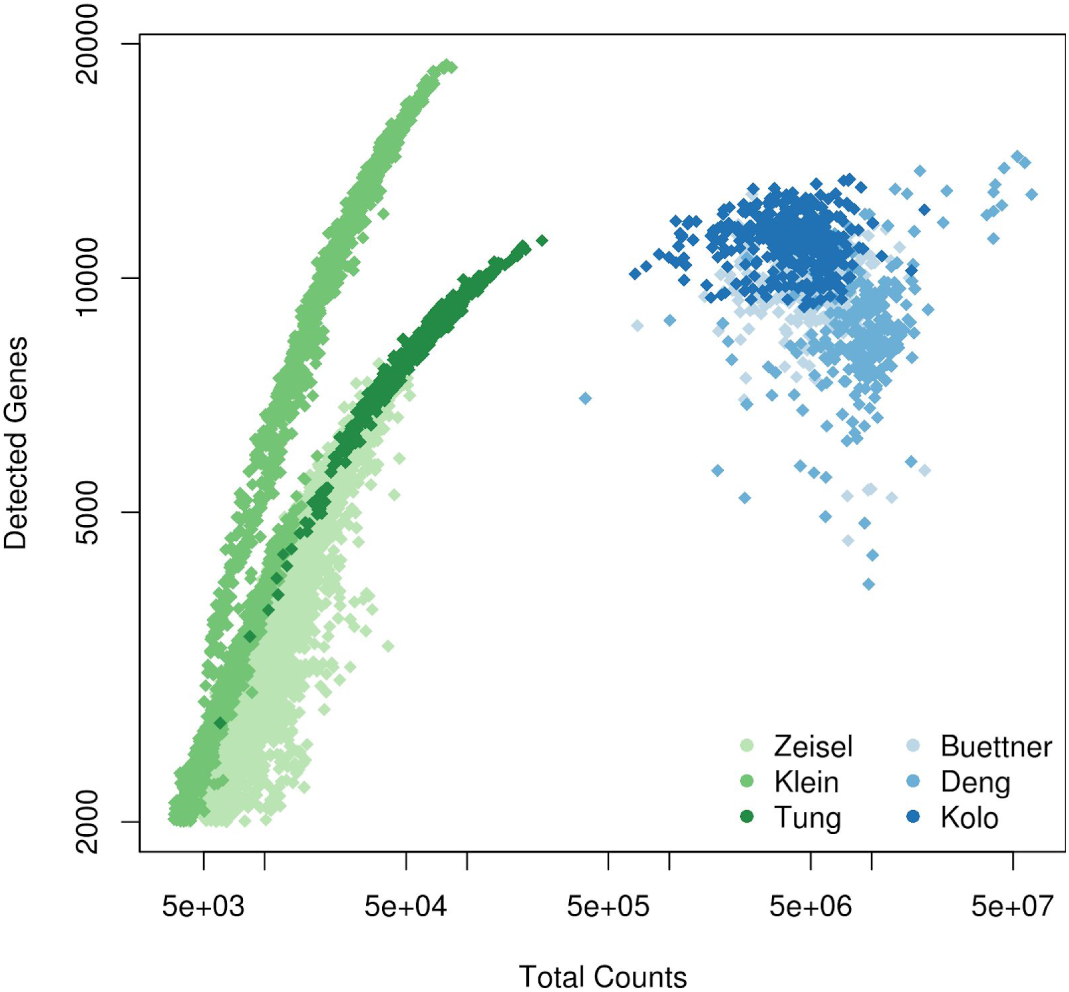
Importance of sequencing depth/tagging efficiency in UMI-tagged (green) vs full-transcript (blue) datasets.

**Figure S3.**
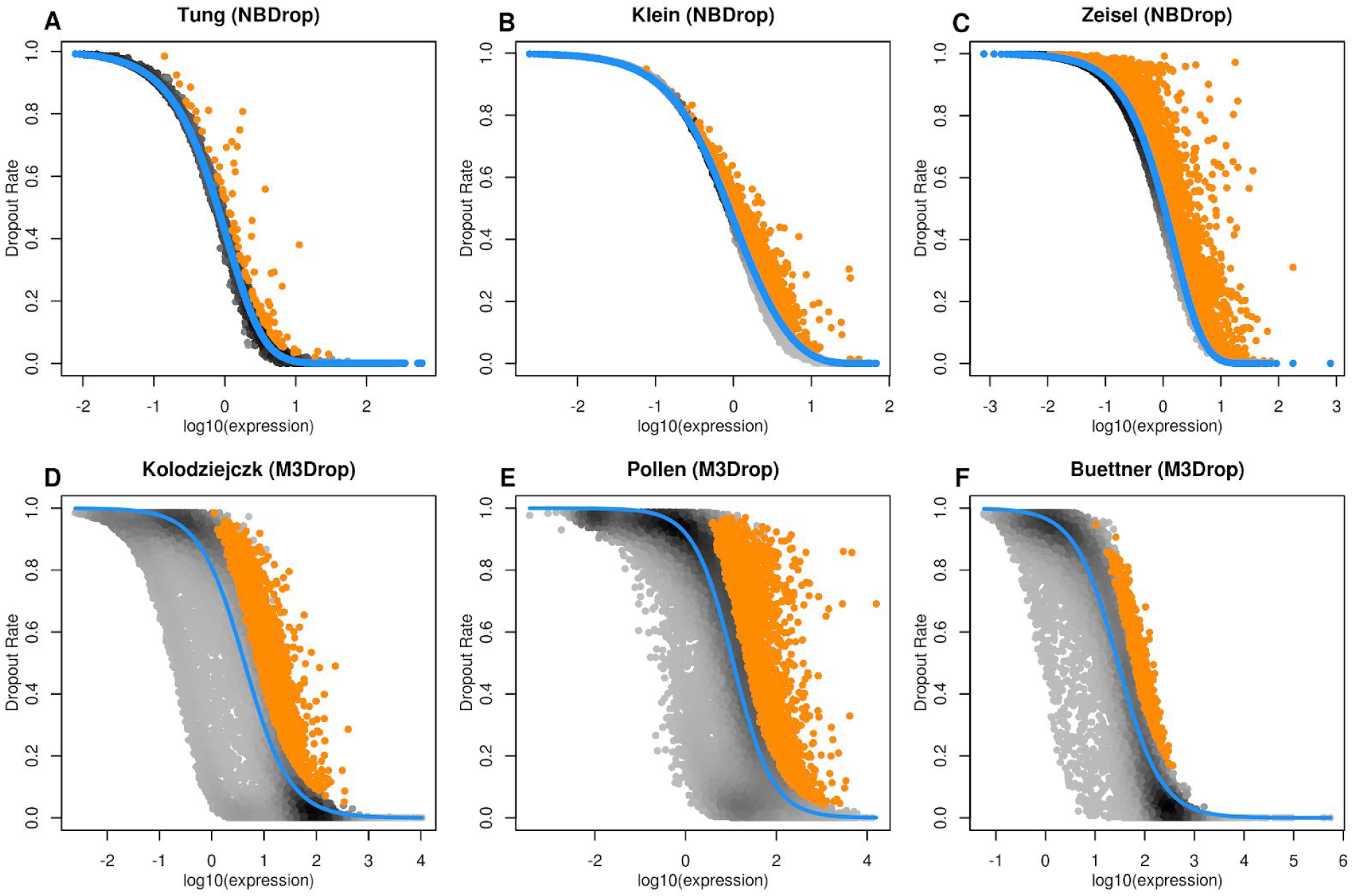
Fitting NBDrop (**A-C**) and M3Drop (**D-F**) to three UMI-tagged and full-transcript scRNASeq datasets respectively. Each point is a gene coloured by the local density of points around it (black = high density). Blue line indicates the fitted relationship between mean and dropout rate from NBDrop or M3Drop respectively. Orange points are significant features using each method.

**Figure S4.**
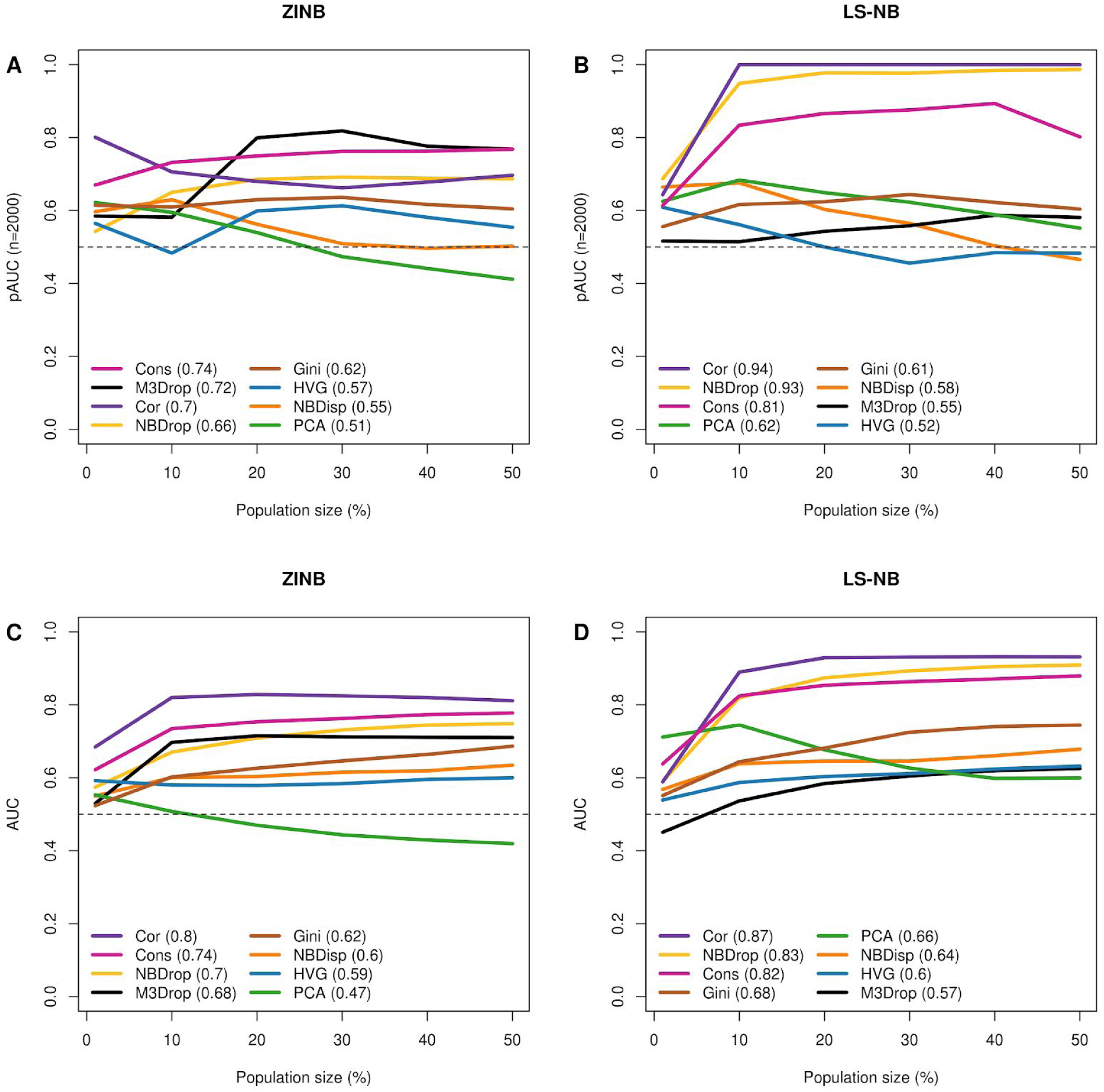
Performance as measured by the AUC for the top 2000 ranked gene (**A,B**) or all genes (**C,D**) when the minor cell-population composes 1, 10, 20, 30, 40 or 50% of the total cells. Count matrices were simulated using a zero-inflated negative binomial (ZINB) or library-size adjusted negative binomial (LS-NB) fit to each of three full transcript and three umi-tagged datasets respectively. Three replicates of 25,000 genes each were performed for each datasets at each subpopulation size. The numbers in the legend represent the average AUC for the six points.

**Figure S5.**
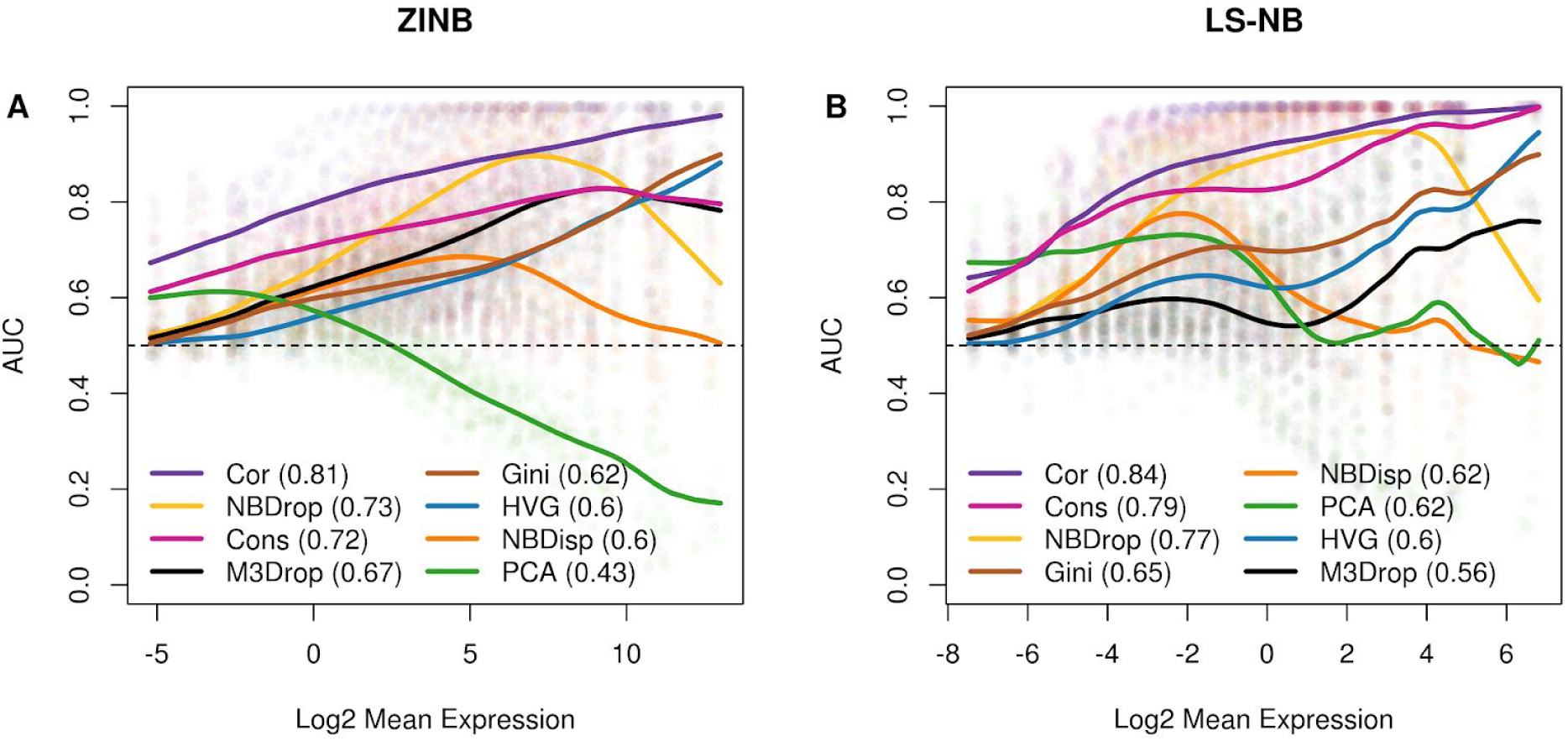
Performance as measured by the AUC for all genes binned into 20 quantiles. Count matrices were simulated using a zero-inflated negative binomial (ZINB) or library-size adjusted negative binomial (LS-NB) fit to each of three full transcript and three umi-tagged datasets respectively. Three replicates of 25,000 genes each were performed for each datasets at each subpopulation size. Points are results for each bin from each replicate, lines are spline-smoothed trends for each feature selection method.

**Figure S6.**
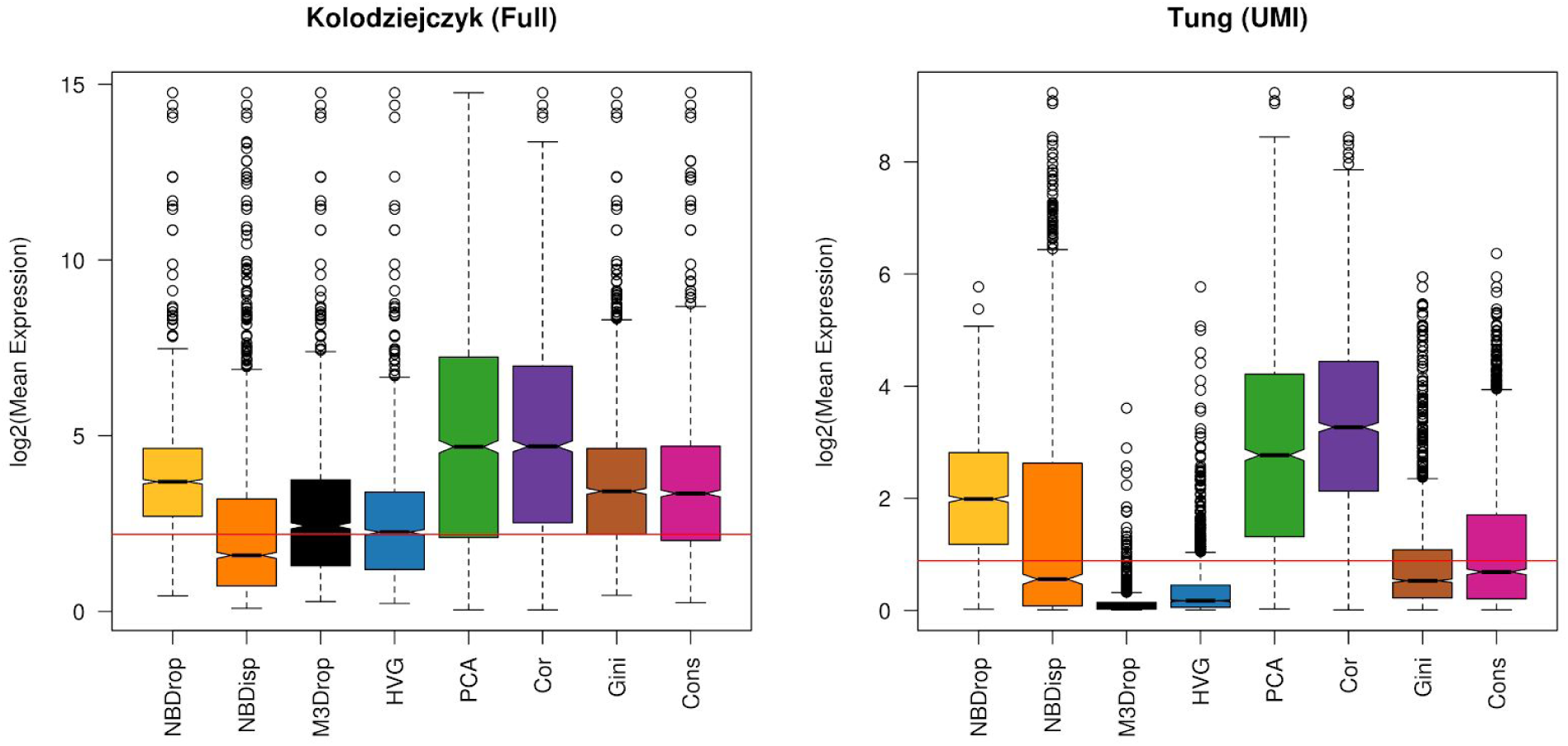
Gene-gene correlations and PCA are biased towards highly expressed genes in real scRNASeq data. Boxplots of the expression level of the top 2000 features identified by each method on the real scRNASeq data. Red line indicates the median expression level for all genes in each dataset.

**Figure S6.**
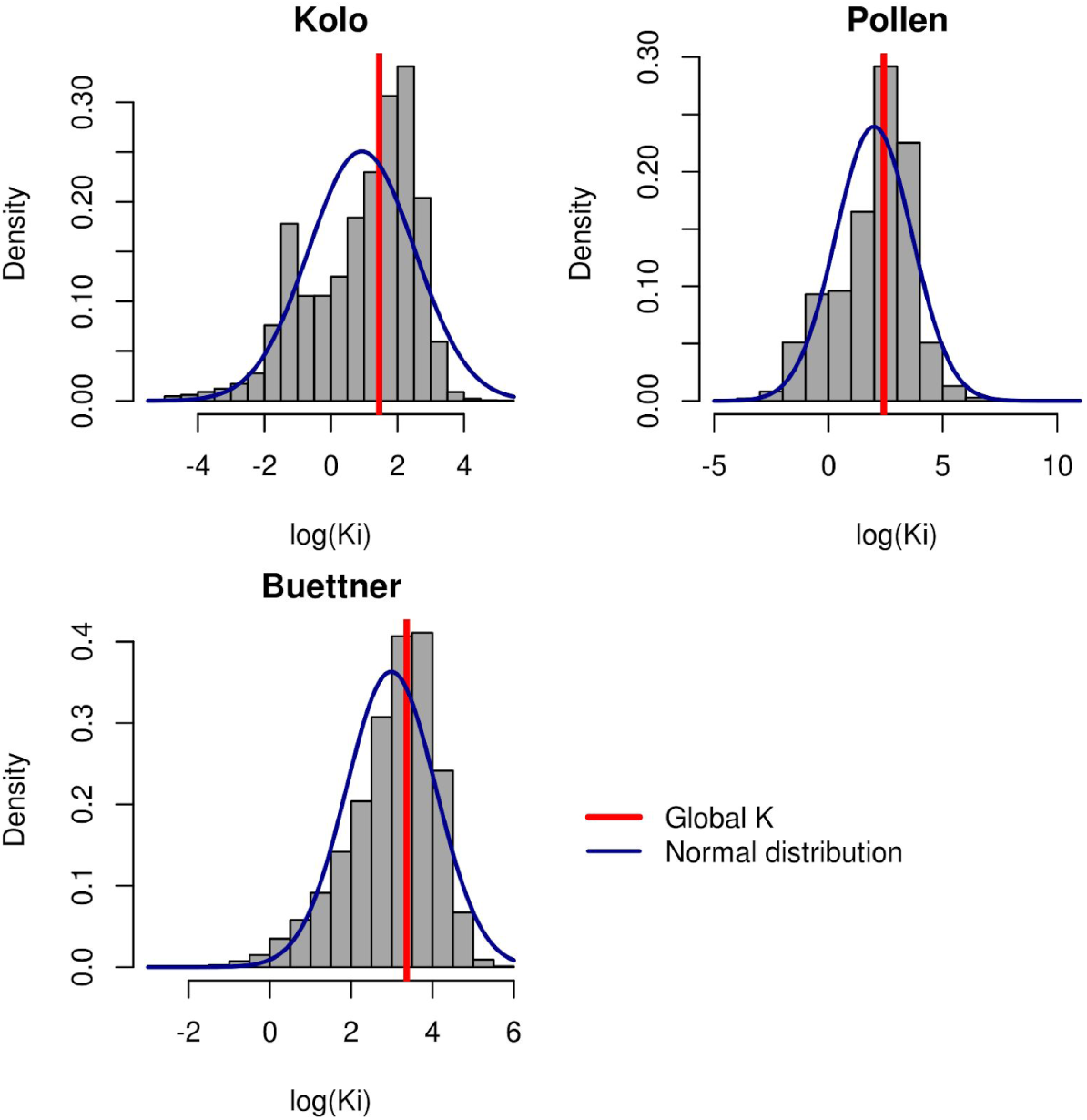
Normal distribution of K_i_ around K_M_

**Figure S7.**
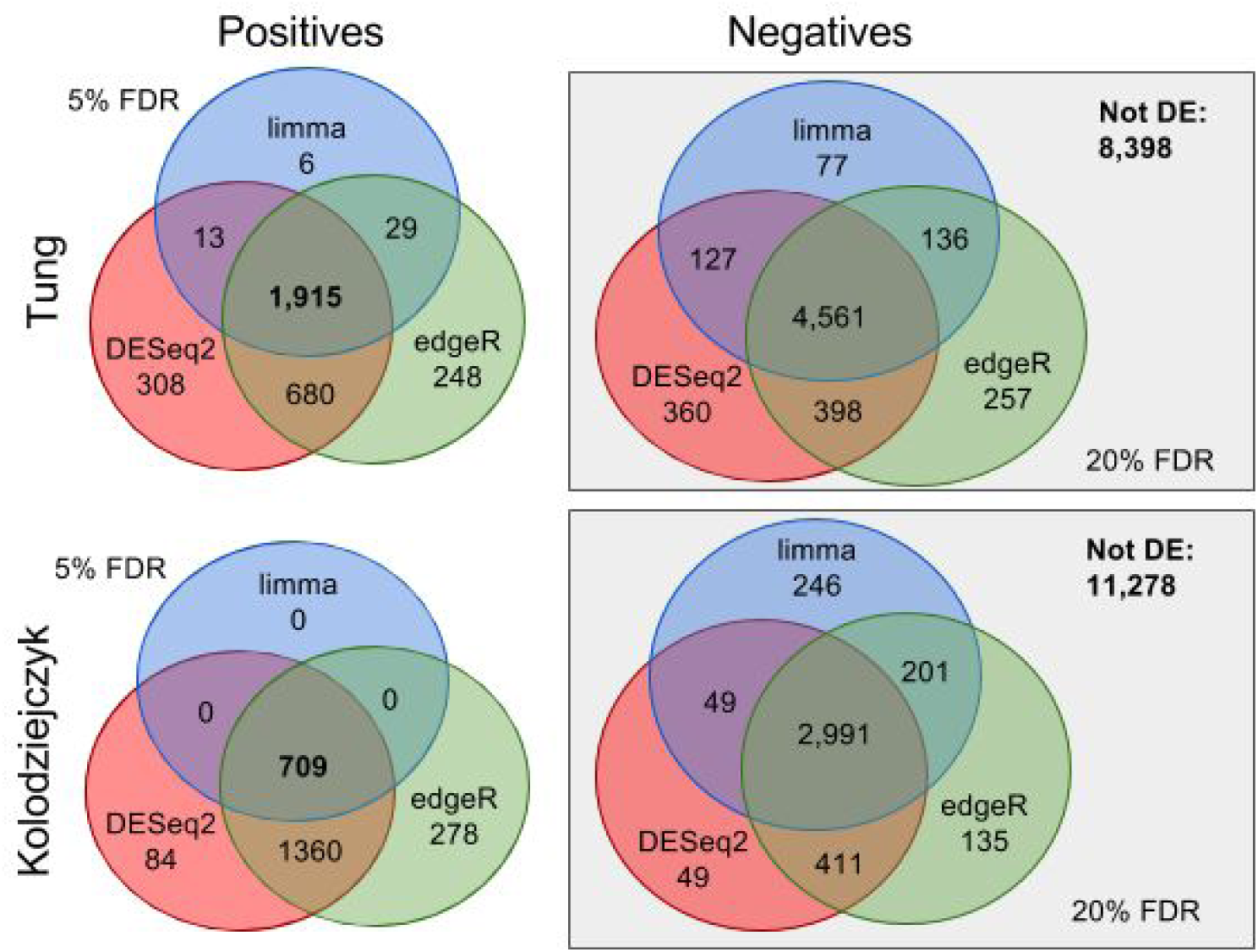
Ground truth DE genes were defined using the intersection (positives) and the complement of the union (negatives) of three standard differential expression method on the respective bulk RNASeq data of the Tung and Kolodziejzck datasets.

**Figure S8.**
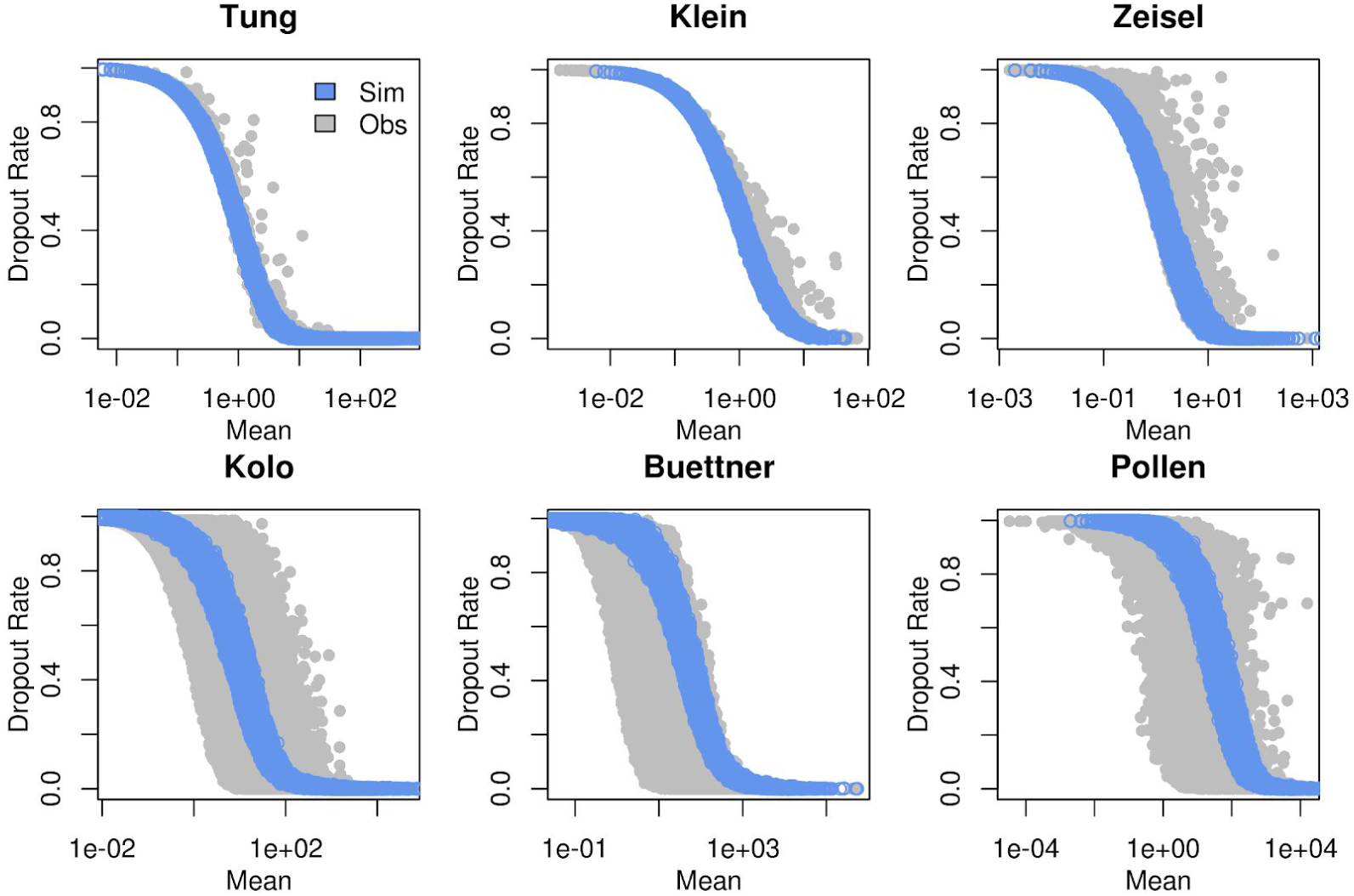
Simulations recapitulate observed relationship between gene expression and dropout rate. Only genes with log fold changes smaller than one (ground truth negatives) are plotted for the simulated data (blue).

**Figure S9.**
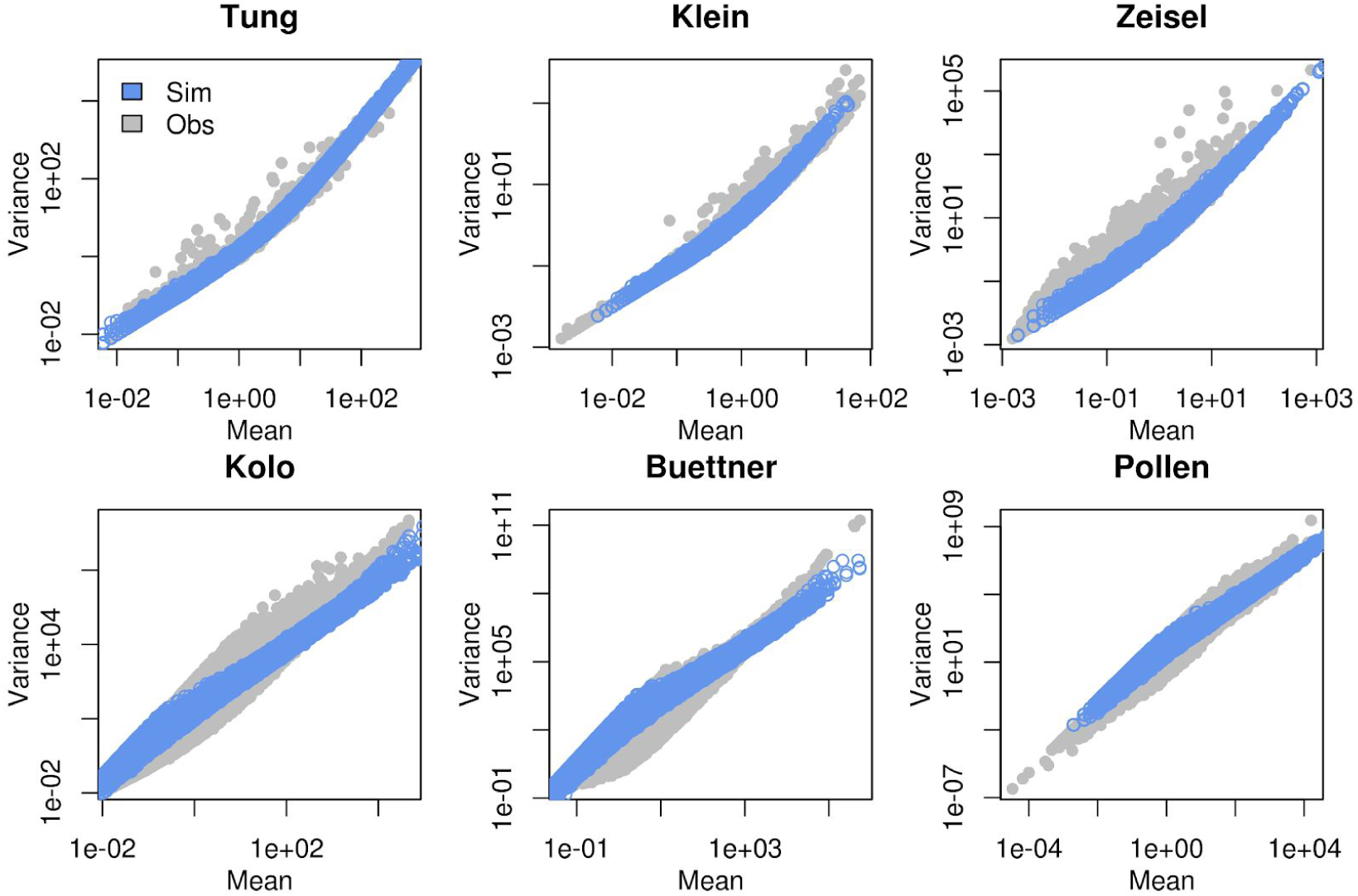
Simulations recapitulate observed relationship between gene expression and variance. Only genes with log fold changes smaller than one (ground truth negatives) are plotted for the simulated data (blue).

